# D1-like receptor activation rescues hippocampal synaptic plasticity and cognitive impairments in the MK-801 schizophrenia model

**DOI:** 10.1101/2024.06.19.599791

**Authors:** Melissa Hernández-Frausto, Emilio J Galván, Carolina López-Rubalcava

## Abstract

Schizophrenia is a disorder with a higher cognitive decline in early adulthood, causing impaired retention of episodic memories. However, the physiological and behavioral functions that underlie cognitive deficits with a potential mechanism to ameliorate and improve cognitive performance are unknown. In this study, we used the MK-801 neurodevelopmental schizophrenia-like model. Rats were divided into two groups: one received MK-801, and the other received saline for five consecutive days (7-11 postnatal days, PND). Using extracellular field recordings in acute hippocampal slices and the Barnes maze task, we evaluated synaptic plasticity late-LTP and spatial memory in freely moving animals in early adolescence and young adulthood. Next, we examined D1-like activation as a mechanism to ameliorate cognitive impairments. Our results suggest that MK-801 neonatal treatment induces impairment in late-LTP expression and deficits in spatial memory retrieval in early adolescence that is maintained until young adulthood. Furthermore, we found that activation of D1-like dopamine receptors ameliorates the impairments and promotes a robust expression of late-LTP and an improved performance in the Barnes maze task, suggesting a novel and potential therapeutic role in treating cognitive impairments in schizophrenia.

**Highlights:**

- MK-801 Schizophrenia model induces impairment in Late-LTP at early adolescence and young developmental stage.
- Barnes maze recall phase is impaired in the MK-801 Schizophrenia model.
- The activation of D1-like receptors promotes recovery and induction of the
- Late-LTP in the MK-801 schizophrenia model in adolescent and young adult rats.
- Activation of D1-like dopamine receptors improves behavioral performance in the MK-801 schizophrenia model in adolescent and young adult rats.

**Graphical Abstract:** 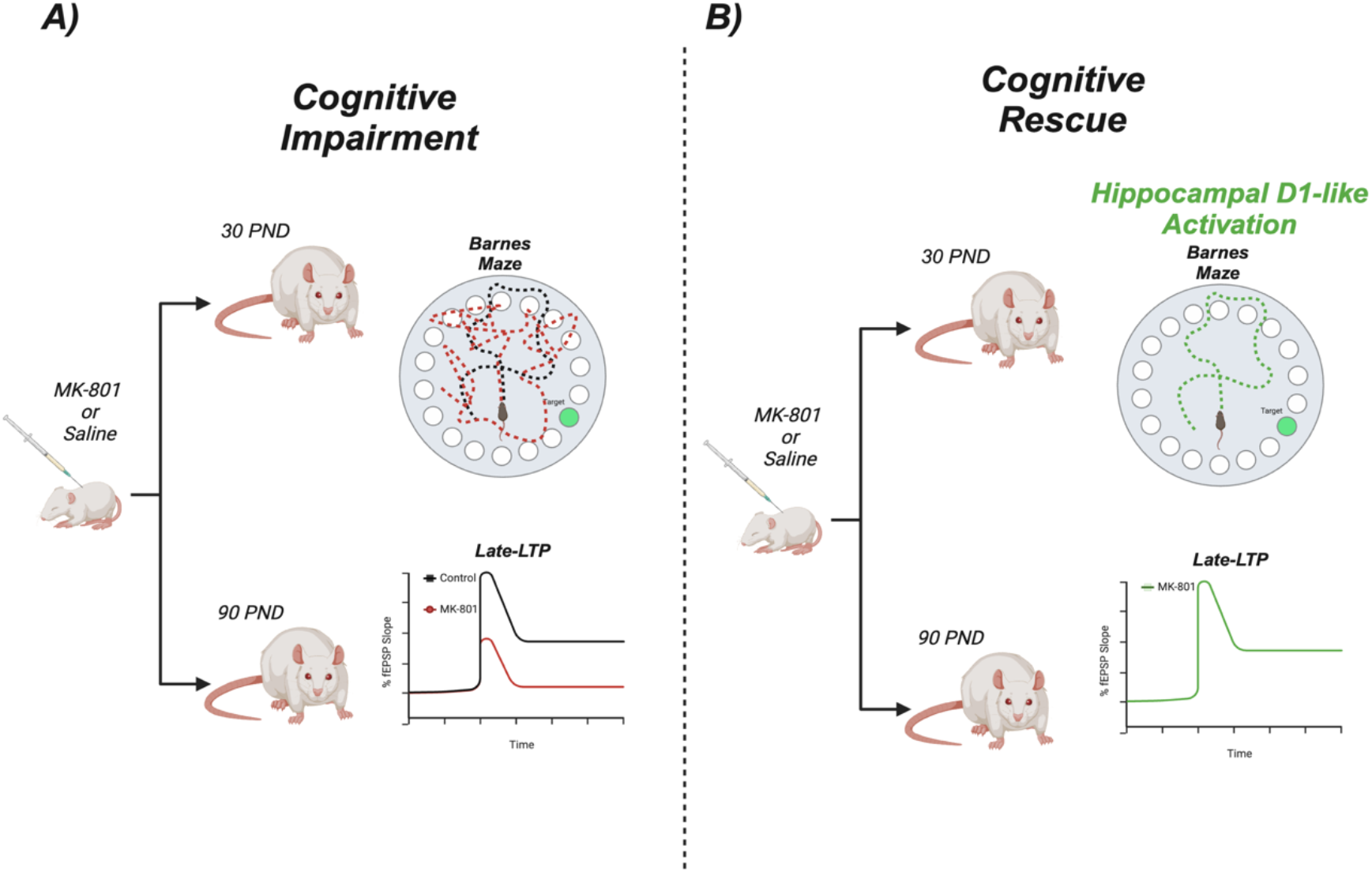

Disturbances in episodic memories are observed in patients with schizophrenia and rodent models. Still, the hippocampal physiological substrates with a potential rescue mechanism related to consolidation of memories have not been elucidated. Here, in vitro electrophysiology and in vivo Barnes maze task are used at two ages in the MK-801 neurodevelopmental schizophrenia-like model. We observed a loss of hippocampal late-LTP and impaired recall phase, and the activation of dopamine D1-like receptors attenuated the impairments with a rescue of both late-LTP and recall phase, suggesting an important role of D1-like receptors activity for episodic memory in schizophrenia.

## 1. Introduction

Schizophrenia is a neuropsychiatric disorder characterized by delusions and hallucinations, with a markedly cognitive decline [1]. The primary cognitive disturbances comprise the acquisition and retention of long-term episodic memories linked to hippocampal function [2], [3], [4]. The symptoms typically manifest during young adulthood with a bare exhibition in the early stages of childhood and young adolescence [2], [5], [6]. Whereas symptoms would not appear until later in the developmental stage, neurodevelopmental impairments are believed to play a critical role in the disease manifestation. An aberrant glutamatergic and dopaminergic transmission maturation has been proposed as a recognized mechanism leading to deficits in the schizophrenia disease course [7], [8].

Regarding cognitive symptoms, hippocampal function disturbances have been reported in patients diagnosed with schizophrenia and in rodent models [9]. Evidence indicates a reorganization of glutamatergic receptors and impairments in synaptic plasticity in the early onset of the disease [9], [10], [11]. In non-pathological conditions, spatial memory consolidation, retention, and retrieval dependence of hippocampal area CA1 function are associated with synaptic plasticity at the late phase of long-term potentiation (late-LTP). Late-LTP promotes the reinforcement of synapses that can last several hours to days [12], [13], [14]. This critical mechanism maintains the synapses’ efficacy and requires synthesizing proteins *de novo* and dopamine as regulators through activating D1-like receptors in glutamate synapses [15], [16], [17], [18]. In the dorsal portion of hippocampal CA1, the most robust dopamine influx comes from the ventral tegmental area (VTA) and locus coeruleus (LC) of dopaminergic and noradrenergic neurons, respectively. These inputs are critical for novelty, as a predictor error, and for long-term memory retrieval [19], [20], [21], [22], [23], [24], [25], [26].

In rodent models, chronic administration of an NMDA receptor antagonist, such as MK-801 (dizocilpine), during neurodevelopment leads to long-term impairments in behavior performance associated with cognitive, negative, and positive-like symptoms and cellular dysfunction such as synaptic plasticity exhibiting a deficiency in the induction of long-term potentiation (LTP) [2], [7], [27], [28]. This NMDA-neurodevelopmental model promotes aberrant connectivity and neurotransmission through the hippocampus. The negative and cognitive impairments in schizophrenia related to hippocampal dysfunction have been extensively described [29], [30]. Recently, we demonstrated the progressive alteration in synaptic efficacy and synaptic plasticity preceding the establishment of behavioral symptoms in a schizophrenia-like model [27]. Moreover, we observed how MK-801 neonatal treatment increased the conductance associated with the firing properties of hippocampal cells in area CA1 [31].

In the present work, we used *in vitro* electrophysiological recordings and *in vivo* freely moving cognitive behavior task, Barnes maze. We aimed to understand the effect of NMDA receptor blockade with MK-801 during the neurodevelopmental period in late-LTP and the Barnes maze task in two ages related to the prodromic and post-dromic manifestation of schizophrenia-like symptoms. Furthermore, our work intends to further extend the restoration of cellular and behavioral impairments through dopaminergic D1-like modulation.

## 2. Experimental procedures

### 2.1. Animals

Animal procedures followed the Mexican Official Norm for the good care and use of laboratory animals “NOM-062-ZOO-1999” and by the local ethics committee of Cinvestav (CICUAL, 0090-14). Female Wistar pregnant rats were obtained from the Pharmacobiology CINVESTAV-SUR vivarium. Female rats and pups were maintained under a 12:12-h light/dark inverted cycle in a controlled temperature at 22 ± 2°C with *ad libitum* access to food and water. Postnatal day (PND) 0 was designated according to the day of birth. Only male pups were used in this study.

### 2.2. MK-801 neonatal treatment

Male pups received subcutaneous injections of MK-801 (0.2 mg/kg) or vehicle saline (control group) from PND 7th to 11th. The pups were maintained under maternal care until weaning at PND 21st. At weanling, the MK-801 and control rats were divided according to age; MK-801 and control groups (30 PND; n= 70) and MK-801 and control groups (90 PND; n= 58). MK-801 was purchased from Sigma-Aldrich (Toluca, Mexico).

### 2.3. Slice preparation and electrophysiological recordings

Electrophysiological and synaptic plasticity recordings at *Schaffer collateral* to CA1 neurons synapse in hippocampal slices from MK-801 and control rats were carried out as described previously [27]. Briefly, rats were anesthetized with pentobarbital (50mg/kg body weight, ip) and decapitated. Brains were exposed and placed in an ice-cold sucrose solution containing (in mM): 210 sucrose, 2.8 KCl, 2 MgSO_4_, 1.25 Na_2_HPO_4_, 26 NaHCO_3_, 6 MgCl_2_, 1 CaCl_2_, and 10 D-glucose, with pH 7.2–7.35, saturated with a 95% O_2_/5% CO_2_ carbogen mixture. Right after, hemispheres were separated through the midsagittal line; tissue was glued to the plate of a VT1000S vibratome (Leica, Nussloch, Germany), and transverse hippocampal slices (385 mm thickness) were obtained. Slices were transferred and maintained for 30min at 33 ± 2 °C in an incubation solution containing (in mM): 125 NaCl, 2.5 KCl, 1.2 Na_2_HPO_4_, 25 NaHCO_3_, 2 MgC_l2_, 1 CaCl_2,_ and 10 glucose, with pH 7.3, continuously bubbled with 95% O_2_/5% CO_2_. After incubation, slices were stabilized at room temperature for at least 60 minutes. For electrophysiological recordings, individual slices were transferred to a submersion recording chamber (submerged in a total volume of 400μl) for at least 20min before the experiments started. Slices were perfused with a standard artificial cerebrospinal fluid (ASCF) solution containing (in mM): 125 NaCl, 3 KCl, 12.5 Na_2_HPO_4_, 25 NaHCO_3_, 2.5 CaCl_2_, 1.5 MgCl_2_, and 10 glucose, maintained at a constant flow (3.5–4 ml/min) at 33 ± 2 °C with an inline solution heater coupled to a temperature controller (TC-324C, Warner Instruments).

Extracellular field excitatory postsynaptic potentials (fEPSP) were evoked with a bipolar electrode and acquired in the *Stratum Radiatum* of area CA1 with a borosilicate glass pipette with a tip resistance of 1–2 MΩ filled with 3M NaCl (Figure 1. B for detailed information). Stimuli were delivered using a high voltage isolator unit (A365D; World Precision Instruments, Sarasota, FL, USA), controlled with a Master-8 pulse generator (AMPI, Jerusalem, Israel) using a stimulation frequency of 0.06Hz, with an intensity range of 15–30μA, and duration of 100 ± 10μs to prevent saturation of responses. Analog responses were amplified with a Dagan BVC-700A amplifier (Minneapolis, MN, USA) coupled to an extracellular head stage (Dagan 8024) and high-pass filtered at 0.3Hz. Responses evoked were displayed on a computer-based scope and digitalized for storage and offline analysis with custom-made software (Lab View system, National Instruments, Austin, TX, USA).

**Figure 1.**
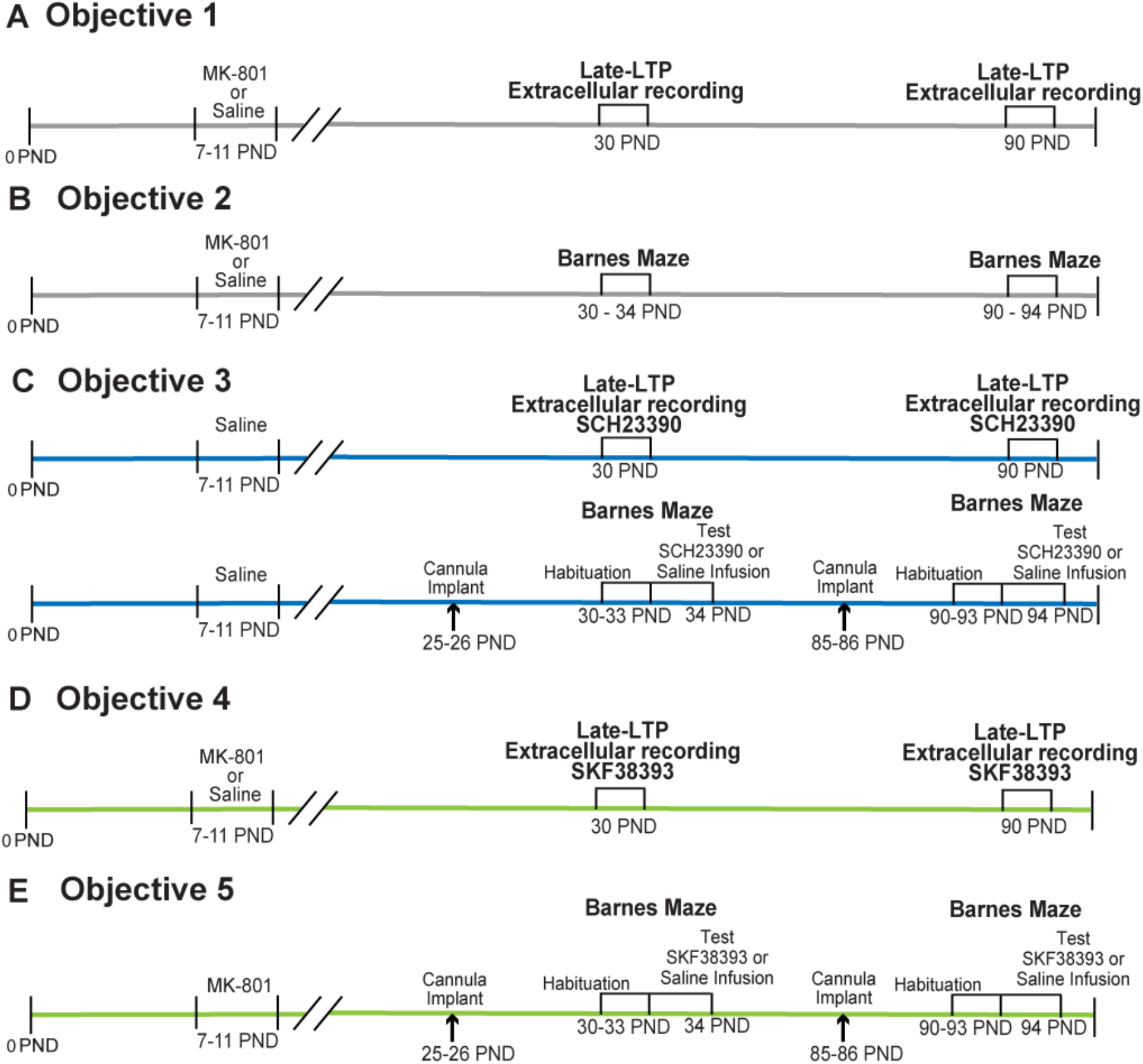
Experimental design of the objectives evaluated. A schematic representation of the timeline is employed to evaluate the neonatal treatment of MK-801 during adolescence and young adulthood, seen as A, B, C, D, and **E.**

To induce the chemical late phase of long-term potentiation (late-LTP), a mixture solution of FSK (forskolin, from Coleus forskohlii, 50µM, Sigma-Aldrich, Toluca Mexico) and IBMX (3-Isobutyl-methylxanthine, 25µM, Sigma-Aldrich, Toluca Mexico) with ACSF was perfused for 25min. This type of late-LTP is NMDA-dependent synaptic plasticity that elicits a PKA-dependent chemical late-LTP maintained for several hours, maximizing the recruitment of several synapses [32], [33], [34]. Increments in the slope of fEPSP were recorded as a % of at least 240min.

For D1-like receptors blockade experiments, a mixture of the D1-like antagonist SCH23390 (R (+)-SCH-23390 hydrochloride at 2µmol, from Sigma-Aldrich, Toluca, Mexico) with ACSF was perfused 20min during baseline (no changes in baseline or fiber volley were observed, data not shown), 25min during FSK-IBMX perfusion, and 15min after FSK-IBMX in acute slices from control animals.

For D1-like receptors activation experiments, a mixture of D1-like agonist SKF-38393 ((±)-SKF-38393 hydrochloride at 5µmol (from Sigma-Aldrich, Toluca, Mexico), with ACSF was perfused 20min during baseline (no changes in fEPSP slope response were observed), 25min during FSK-IBMX perfusion, and 15min after FSK-IBMX in acute slices from MK-801 animals. D1-like receptor agonist and antagonist doses were chosen as similar to previous reports, and we never observed changes in basal fEPSP slope or epileptiform activity in the brain slices [20], [26], [35], [36], [37], [38], [39], [40].

### 2.4. Surgical procedure

For *in vivo* experiments, rats were implanted unilaterally with a guide cannula 1mm above the *Stratum Radiatum* of dorsal CA1 as follows: in mm, + 2.5 anteroposterior (AP), 1.2 mediolateral (ML), and 1.0 dorsoventral (DV); all coordinates relative to bregma (unilateral cannula implants were prepared to prevent hyperlocomotion related to dopamine D1-like receptors effects that may unmask the fundamental cognitive role).

The effects of D1-like antagonist and agonist were tested using SCH23390 and SKF38393, respectively (Bach et al., 1999; Broussard et al., 2016; Huang & Kandel, 2006; Lemon & Manahan-Vaughan, 2006; Morris et al., 2003; Wiescholleck & Manahan-Vaughan, 2014). Control groups with local infusion of saline solution were also included in all the experiments, and rats were given four days to recover before the experiments started.

### 2.5. Barnes Maze test

The Barnes Maze is a circular black platform with 18 evenly spaced holes around its edge. It evaluates a rat’s ability to learn and remember spatial functions associated with hippocampal spatial functioning [41]. Two weeks before the behavioral test, rats were habituated to the experimenter user and the experimentation room to avoid anxiogenic behavior that may unmask the actual cognitive assessment.

The behavioral task is divided into habituation and retention phases, with a total duration of five days. A concealed black chamber is situated beneath one of the holes, and visual cues are distributed throughout the maze. Mild aversive stimuli in the form of bright lights and white noise are also present. The test was consistently performed at the same daily time (involving a dark cycle of the rats). The escape box was maintained at the same location as the spatial cues, whereas the holes were constantly moved and cleaned before each trial to avoid odor cues.

#### Habituation phase

Rodents underwent a training process called the habituation phase. Here, rats were exposed to the escape chamber and maze arena twice daily, with a 15-minute break. During this habituation phase, rats were familiarized with the target chamber for one minute and then allowed to explore the arena for three minutes. If the animal failed to reach the target chamber within this period, it was gently guided into the box and left for one minute of rest.

#### Retention phase

Twenty-four hours after the last habituation trial, the retention phase was conducted to evaluate memory retrieval and ended once the rat reached the escape chamber or three minutes had elapsed.

All trials were recorded and analyzed using Any-maze software to determine the total latency and total number of errors.

##### For unilateral local infusion in dorsal CA1 experiments

First, a D1-like antagonist (SCH23390) at a concentration of 2µmol was locally infused to control groups of 30 and 90 PND to assess the involvement of D1-like receptors in the memory retention phase. Next, a D1-like agonist (SKF38393) at a concentration of 5µmol was locally infused to MK-801 groups of 30 and 90 PND in independent groups. Concentration infusions were based on previous work assessing the role of D1-like receptors’ involvement with *in vivo* cognitive behaviors and did not observe any stereotyped behaviors associated with hyperdopaminergic activity [20], [26], [35], [36], [37], [38], [39], [40].

### 2.6. Statistical Analyses

Group measures are expressed as the mean ± standard error of the mean (SEM). Normality was tested with the Shapiro-Wilk test, and equal variance was evaluated before the ANOVA tests. A t-test was used for the retention phase during the Barnes maze. A two-way ANOVA followed by Holm-Sidak was used to compare D1 and D4 and for group comparisons in the Barnes maze task. For electrophysiology analysis, changes in the CA1 fEPSP slope were determined by comparing the mean of fEPSP during FSK-IBMX perfusion, 120min, and 240min after perfusion within baseline by a t-test.

### 2.7. Experimental Design

A total of 128 rats were used in the present study. For the first set of experiments, male rats were divided according to age, at 30 PND and 90 PND, and according to group MK-801 and control group. Afterward, they were divided according to objectives as follows (Figure 1A-E):

#### Objective 1

To address that late-LTP in the hippocampal CA1 area is impaired in the MK-801 schizophrenia-like model in adolescence and young adulthood. One slice per rat in each condition was occupied. The electrophysiological recordings were made as follows: at 30 PND, 12 rats were used for control and 11 for the MK-801 group (a total of 23). At 90 PND, 7 rats were used for the control group and 7 for the MK-801 group (a total of 14 rats) (Figure 1, panel A).

#### Objective 2

To determine that neonatal treatment with MK-801 impairs spatial memory in the adolescent and adult developmental stages in the Barnes maze test. At 30 PND, for the behavioral assay with the Barnes maze, 15 rats were used as a control group and 12 for the MK-801 group (a total of 27 rats). At 90 PND, 15 rats were used for the control group and 13 for the MK-801 group (a total of 28 rats) (Figure 1, panel B).

#### Objective 3

To investigate if blockade of dopamine D1-like receptor impairs late-LTP in CA1 and decreases performance during a spatial memory task in control conditions from adolescent and adult animals. For late-LTP electrophysiological recordings, one slice per control group rat was occupied. Bath perfusion of the antagonist of D1-like, SCH23390, for 60 minutes was performed (no changes at the baseline were observed, data not shown). At 30 PND, 5 rats (1 slice per rat) and 6 rats (1 slice per rat) at 90 PND were employed. For the Barnes maze task, local infusion of the antagonist of D1-like SCH23390 or saline only was done. At 30 PND, 7 rats were locally infused only with saline and 6 with SCH23390 (a total of 13 rats). At 90PND, 7 rats were locally infused only with saline and 8 with SCH23390 (a total of 15 rats) (Figure 1, panel C).

#### Objective 4

To investigate if the activation of the D1-like receptor reverts MK-801 neonatal treatment impairments and promotes late-LTP at adolescent and adult developmental stages. For the electrophysiological recordings, 1 slice per rat of MK-801 at 30PND and 90 PND groups were occupied. Bath perfusion of the agonist of D1-like, SKF38393, for 60 minutes was performed. At 30 PND, 10 rats (1 slice per rat) and 6 rats (1 slice per rat) at 90 PND were employed (Figure 1, panel D).

#### Objective 5

*T*o evaluate if the local infusion of SKF38393 in the dorsal hippocampal CA1 area ameliorates retention phase impairment at adolescent and adulthood stages in MK-801 treated animals in the Barnes maze test. For the behavioral assay, local infusion of the agonist of D1-like SKF38393 or saline only was performed. At a rate of 0.25μmol per minute for 5 minutes (total concentration of 5μmol) with 5 minutes of rest before the behavioral test. At 30 PND, 8 rats were locally infused only with saline and 11 with SKF39303 (a total of 19 rats). At 90PND, 12 rats were locally infused only with saline and 14 with SCH23390 (a total of 26 rats) (Figure 1, panel E).

## 3. Results

### 3.1. Late-LTP in the hippocampal CA1 area is impaired in the MK-801 schizophrenia-like model in adolescence and young adulthood

Late-LTP is considered the cellular mechanism for long-term memory consolidation and retrieval in the hippocampal area CA1 [42], [43]. Therefore, we explored if late-LTP is impaired in early adolescence (30 PND) and remains until young adulthood (90 PND) in the MK-801 schizophrenia-like model (see methods and schematic from Figure 1, panel A and Figure 2, panel A).

**Figure 2.**
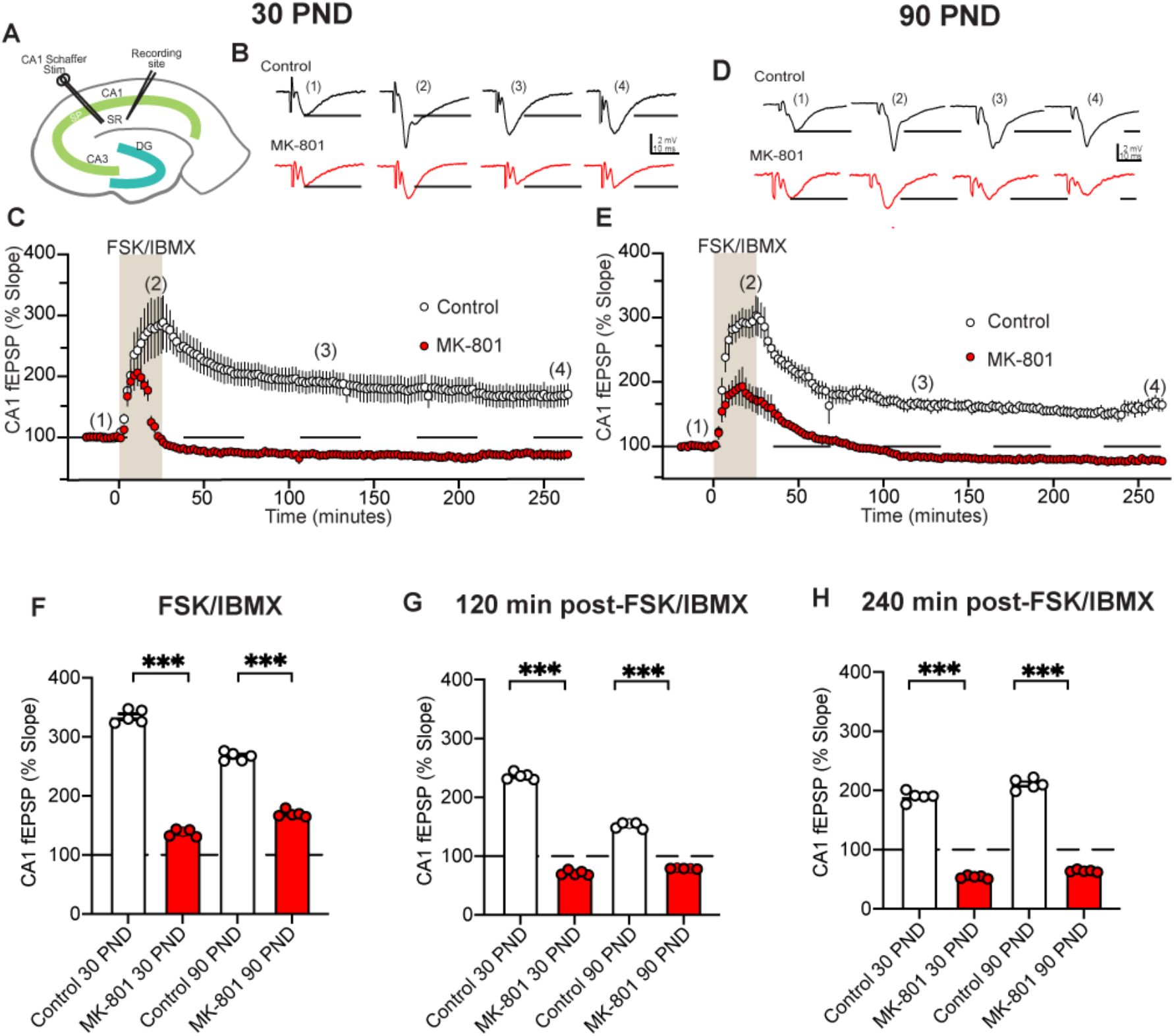
MK-801 neonatal treatment impairs late LTP at 30 and 90 PND. **A)** Location of stimulation and recording electrodes in *Stratum radiatum* (SR) from dorsal CA1 area. **B)** Representative traces of CA1 fEPSP at 30 PND. **C)** Time-course graph of synaptic response in control (empty symbols) and MK-801 (red symbols) conditions. **D)** Representative traces at 90 PND. **E)** Time-course graph of synaptic response in control (empty symbols) and MK-801 (red symbols) conditions. **F)** Bar graph with an average of FSK-IBMX perfusion at 30 PND and 90 PND control and MK-801 treated groups. **G)** Bar graph with an average of fEPSP at 120min at 30 PND and 90 PND control and MK-801 treated groups. **H)** Bar graph with an average of 240min at 30 PND and 90 PND control and MK-801 treated groups.

First, we explored if the FSK-IBMX mixture can induce late-LTP at 30 PND in the MK-801 schizophrenia-like model. In extracellular field recordings (fEPSP) in the dorsal CA1, we found that FSK-IBMX induced a transient increase in the fEPSP slope response. However, this response decreased and remained below basal levels for the 240-minute recording period, unlike control conditions, which showed a robust increase in the fEPSP slope response (Figure 2B-C). Next, at 90 PND, we observed that the FSK-IBMX mixture caused a temporary increase in the fEPSP slope response that decayed under basal levels and lasted for 240min of recording. In contrast, we observed an increased fEPSP slope in control conditions and maintained during 240min of recording (Figure 2D-E).

By comparing the fEPSP slope from the last 5min of the perfusion of the FSK-IBMX mixture at both ages, we observed an increase in the fEPSP slope of slices from MK-801 animals; however the increment was lower compared to control conditions at 30 and 90 PND (at 30 PND t= 38.087, p<0.001 Control Vs MK-801; At 90 PND t= 0.001 Control Vs MK-801, p=1; see Figure 2F). Furthermore, we paralleled the fEPSP slope at 120min post-FSK-IBMX perfusion. At 30 and 90 PND, there is a consistent depression of the fEPSP response, whereas in control conditions, it maintains above the basal levels (At 30 PND t= 40.805, p<0.001 Control Vs MK-801; At 90 PND t= 14.503, p<0.001 Control Vs MK-801; Figure 2G).

Lastly, when comparing the fEPSP slope at 240min after FSK-IBMX perfusion, slices from MK-801 animals fail to recover to basal conditions, whereas, in controls, there is a consistent increase in fEPSP slope response at 30 and 90 PND (At 30 PND t= 32.763, p<0.001 Control Vs MK-801; At 90 PND t= 35.350, p<0.001 Control Vs MK-801; Figure 2H). Our data indicates that in the MK-801 schizophrenia-like model, there is an impairment in late-LTP as early as 30 PND, persisting until 90 PNDs. In contrast, in control conditions, FSK-IBMX can induce a robust late-LTP potentiation that persists during the 240 minutes of recording.

### 3.2. Neonatal treatment with MK-801 impairs spatial memory in the adolescent and adult developmental stages in the Barnes maze test

Barnes maze test is a trial widely used to study spatial memory and is highly sensitive to hippocampal function and dysfunction in neurological disease rodent models [44], [45]. Therefore, we evaluated how the MK-801 schizophrenia-like model modifies the behavioral response during the Barnes maze at early adolescence and young adulthood periods (30 and 90PND, respectively; see Figure 1, panel B and Figure 3, panel A for experimental design details).

**Figure 3.**
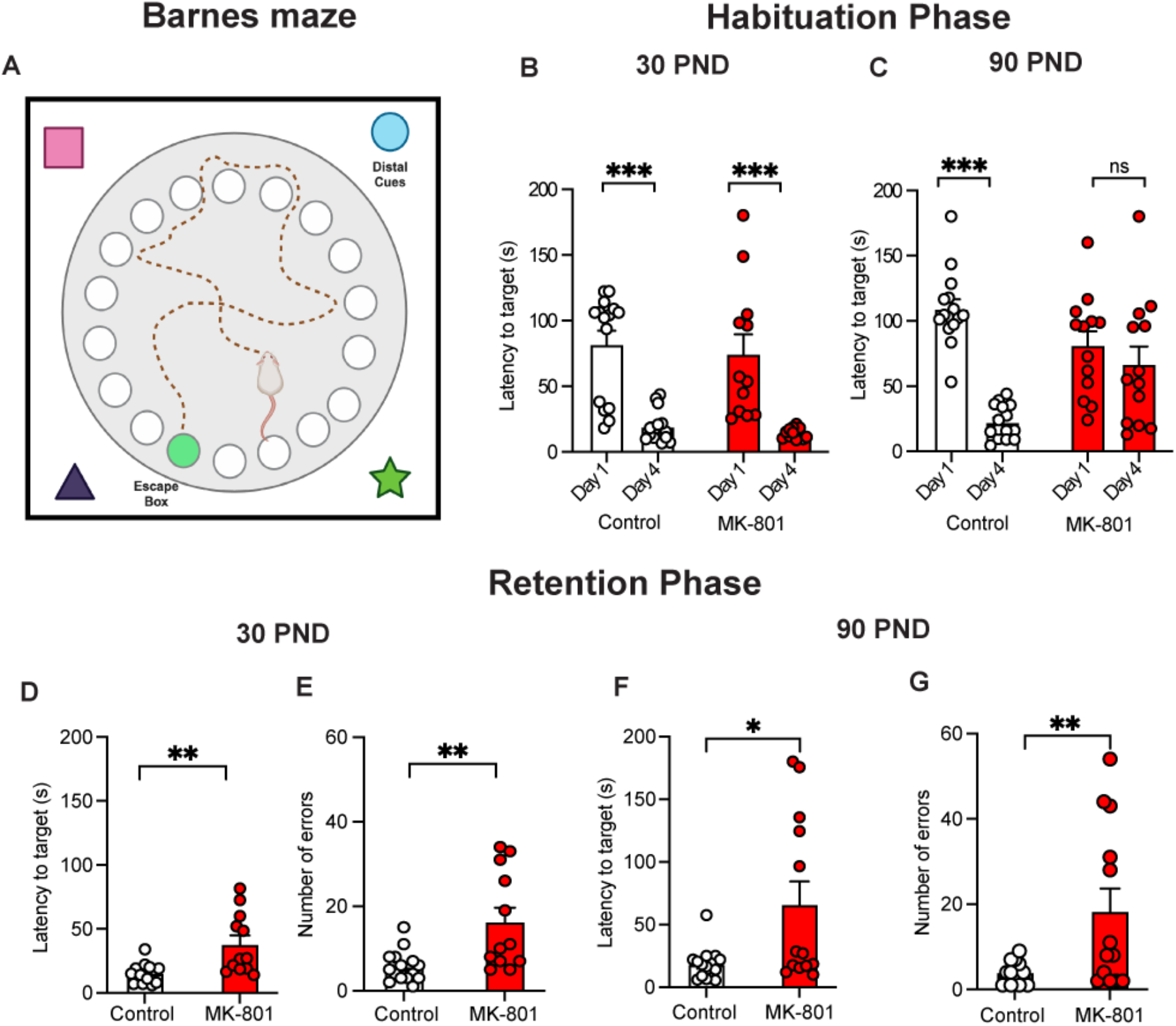
MK-801 neonatal treatment impairs the spatial memory retention phase in the Barnes maze task at 30 and 90 PND. **A)** Representation of Barnes maze task (created with BioRender). **B)** Latency of day 1 (D1) and day 4 (D4) in control (empty symbols) and MK-801 (red symbols) groups at 30 PND. **C)** Latency of day 1 (D1) and day 4 (D4) in control (empty symbols) and MK-801 (red symbols) groups at 90 PND. **D)** At 30 PND, latency on the 5^th^ day in control (empty symbols) and MK-801 group (n = red symbols). **E)** At 30 PND, the number of errors on the 5^th^ day in control (empty symbols) and MK-801 group (red symbols). **F)** At 90 PND, latency on the 5^th^ day in control (empty symbols) and MK-801 group (red symbols). **G)** At 90 PND, the number of errors on the 5^th^ day in control (empty symbols) and MK-801 group (red symbols).

During the acquisition phase at 30 PND, animals treated with MK-801 showed decreased latency to find the target hole and enter the goal chamber on day four compared to day 1 (t= 4.918, p<0.001, Figure 3B), similar to control conditions (t= 4.923, p<0.001, Figure 3B). No differences were observed between groups (Day 1, t= 0.255, p= 0.800, Control Vs. MK-801; Day 4, t= 0.433, p= 0.667). Nonetheless, at 90 PND, animals treated with MK-801 cannot learn and find the target goal and enter the goal chamber. On day 4, they show similar latency as on day 1 (t= 1.085, p=0.283, Figure 3C). In contrast, the control group exhibits a decreased latency to find the target hole and enter the goal chamber on day 4 compared to day 1 (t= 7.022, p<0.001, Figure 3C).

In the retention phase at 30 and 90 PND, at 30 PND, the MK-801 treated group had longer latency (t= 3.513, p<0.01, Figure 3D) and a more significant number of errors (t= 3.353, p<0.01, Figure 3E) in finding the target hole compared to the control group. Similarly, at 90 PND, the MK-801 treated group has an increased latency (t= 2.739, p<0.05, Figure 3F) and a higher number of errors (t= 2.910, p<0.01, Figure 3G) to find the target hole compared to the control group. This suggests that MK-801 treatment, as a schizophrenia-like model, impairs retention and spatial memory processing since early adolescence and worsens during young adulthood.

### 3.3. Blockade of dopamine D1-like receptor impairs late-LTP in CA1 and decreases performance during a spatial memory task in control conditions from adolescent and adult animals

Neuromodulation of dopamine is important for hippocampal cognitive function and episodic memory consolidation and retrieval. Blockade of dopamine function via D1-like receptors decreases memory and synaptic plasticity in late-LTP [35], [46], [47], [48]. Therefore, we aimed to investigate whether blocking D1-like receptors during the retention phase under normal conditions could simulate the cognitive deficits seen in MK-801-treated animals. Consequently, we administered a D1-like antagonist (SCH23390) and evaluated its impact on the mechanisms associated with spatial memory retrieval, both *in vitro* (late-LTP) and *in vivo* (test phase of Barnes maze). As in our previous study, we separated the animals into two age groups: 30 and 90 PND. We then conducted independent evaluations of electrophysiological and behavioral experiments, as described in the methods and experimental design section of Figure 1, panel C. To assess if blockade of D1-like receptors impairs late-LTP at 30 PND. First, we observed that in the perfusion of FSK-IBMX in the presence of SCH23390, there is a transient increase in the fEPSP slope that decreases and remains below the baseline during the 240min of recording (Figure 4A-B). Similarly, at 90 PND, FSK-IBMX perfusion in the presence of SCH23390 induces a transient increase of the fEPSP slope, decays under basal levels, and is maintained below the baseline during the 240min of recording (Figure 4C-D). Additionally, we compared the fEPSP slope from the last 5min of FSK-IBMX and SCH23390 perfusion in both age groups. We observed a transient increase in the fEPSP slope at 30 and 90 PND compared to their baseline (At 30PND, t=28.59, p<0.001; at 90 PND, t= 71.00, p<0.001; Figure 4E). After 120min of FSK-IBMX and SCH23390 perfusion, we observed a decay in the fEPSP slope at 30 and 90 PND compared to their baseline (At 30PND, t=4.18, p<0.0013; at 90 PND, t= 11.59, p<0.001; Figure 4F). Lastly, at 240min after the mixture of FSK-IBMX with SCH23390, the fEPSP slope was not different from baseline conditions (At 30PND, t=0.43, p<0.67; at 90 PND, t= 0.18, p<0.85; Figure 4G).

**Figure 4.**
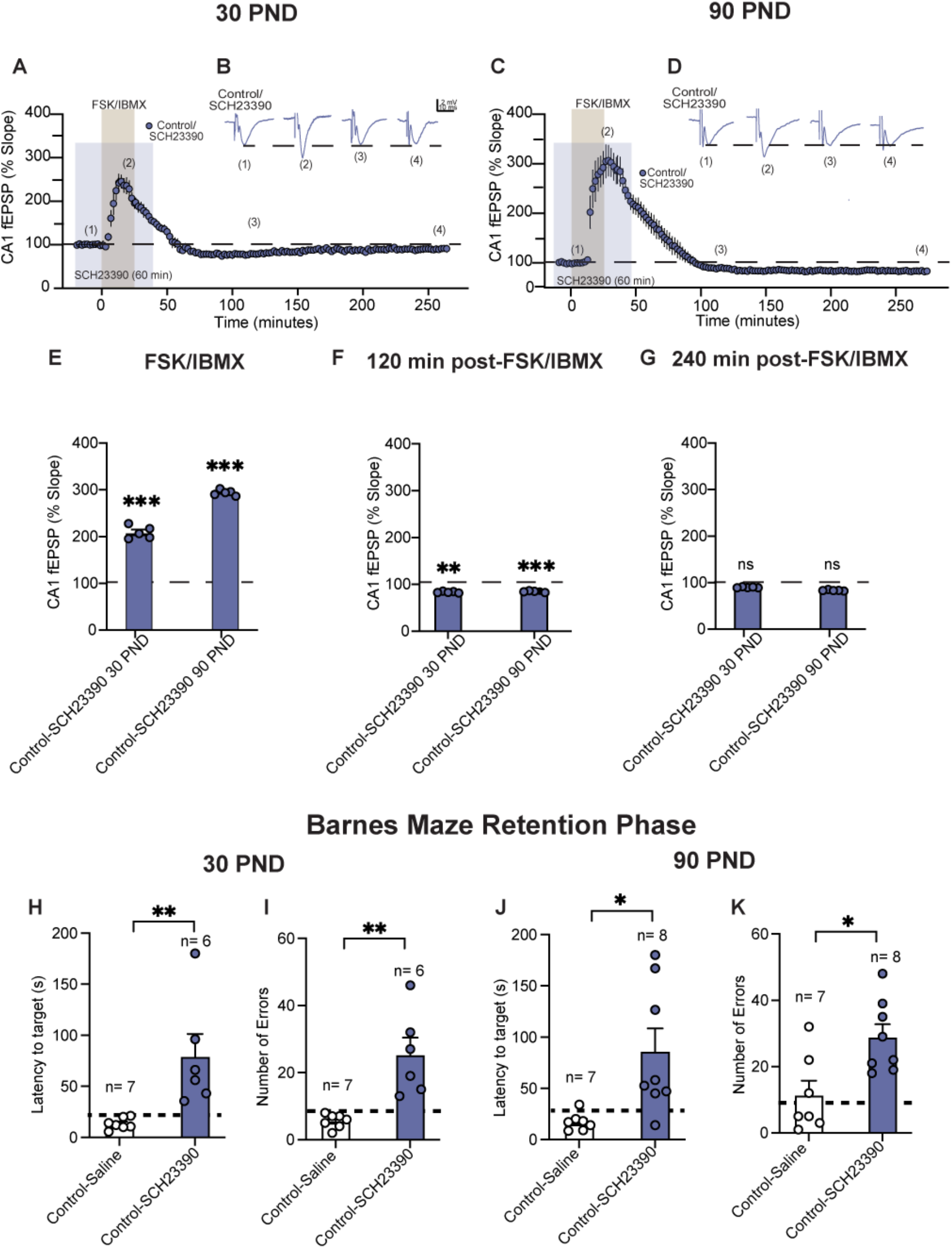
SCH23390 (D1-like antagonist) impairs late-LTP and disrupts the spatial memory retention phase in control conditions at 30 and 90 PND. **A)** Time course graph of synaptic response during SCH23390 conditions (blue symbols). **B)** Representative traces at 30 PND. **C)** Time course graph of the synaptic response of 90 PND animals during SCH23390 conditions (blue symbols). **D)** Representative traces at 90 PND. **E)** Bar graph of fEPSP slope changes in response to FSK-IBMX in 30 PND and 90 PND control-SCH23390 groups. **F)** Bar graph of fEPSP slope changes at 120min in 30 PND and 90 PND control-SCH23390 groups. **G)** Bar graph of fEPSP slope changes at 240min in 30 PND and 90 PND control-SCH23390 groups. **H)** At 30 PND, latency on the 5^th^ day in control-saline (empty symbols) and control-SCH23390 groups (blue symbols). **I)** At 30 PND, the number of errors on the 5^th^ day in control-saline (empty symbols) and control-SCH23390 groups (blue symbols). **J)** At 90 PND, latency on the 5^th^ day in control-saline (empty symbols) and control-SCH23390 groups (blue symbols). **K)** At 90 PND, the number of errors on the 5^th^ day in control-saline (empty symbols) and control-SCH23390 groups (blue symbols).

In an independent set of experiments, to determine if similar results were observed in control groups *in vivo*. We tested the effect of the local infusion of a D1-like antagonist (SCH23390) or saline only into dorsal CA1 during the retention phase of the Barnes maze at 30 PND and 90 PND. The experimental design is described in Figure 1, panel C).

Local unilateral infusion of SCH23390 in dorsal CA1 at 30 PND resulted in higher latency (t= 3.267, p<0.0075; Figure 4H) and increased number of errors (t= 4.168, p=0.0016; Figure 4I) to locate the goal chamber when compared to the saline-control group. Similar to our previous results, at 90 PND, the control-SCH23390 group exhibited an increased latency (t=2.886, p<0.0127; Figure 4J) and a more significant number of errors (t=3.012, p<0.0100; Figure 4LK) to locate the target hole compared to the control-saline group.

With these findings, our results suggest that in control conditions, dopamine through D1-like receptors facilitates late-LTP and spatial retrieval mechanisms in the Barnes maze, similar to our findings with the MK-801 schizophrenia-like model.

### 3.4. Activation of the D1-like receptor reverts MK-801 neonatal treatment impairments and promotes late-LTP at adolescent and adult developmental stages

Our previous findings suggest that blockade of D1-like receptors impairs late-LTP, similar to our MK-801 results. We aimed to determine the role of activation of D1-like receptor with SKF38393 agonist in late-LTP at 30 and 90 PND in MK-801-treated animals. Therefore, hippocampal slices from 30 and 90 PND MK-801 neonatal treated animals were prepared for extracellular field recordings (see methods, experimental design in panel D figure 1 and schematic from Figure 2 panel A for details).

At 30 PND, perfusion of FSK-IBMX in the presence of SKF38393 leads to an increase in the fEPSP slope of the CA1 response that remains above baseline for 240min, resulting in consistent late-LTP synaptic plasticity (Figure 5A-B), compared to baseline. At 90 PND, the perfusion of FSK-IBMX in the presence of SKF38393 raises the fEPSP slope compared to the baseline and maintains over the baseline during the 240min of recording with a prominent late-LTP (Figure 5C-D).

**Figure 5.**
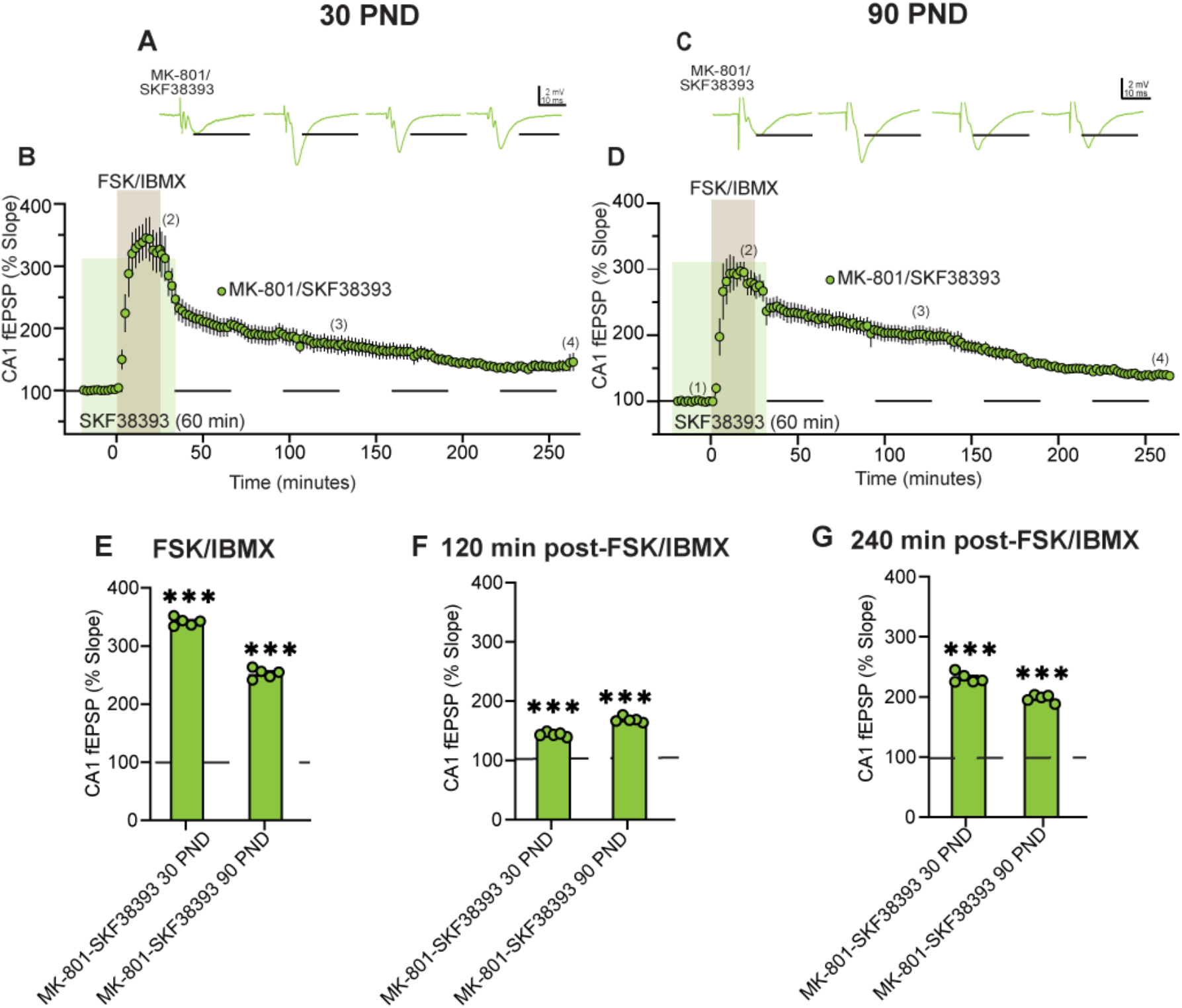
Activation of the D1-like receptor (D1/ D5) reverts MK-801 treatment impairments and promotes the late phase of LTP during adolescence and young adulthood. **A)** Representative traces at 30 PND. **B)** Time-course graph of synaptic response during SKF38393 (green symbols) at 30PND. **C)** Representative traces at 90PND. **D)** Time-course graph of synaptic response during SKF38393 (green symbols) at 90 PND. **E)** Bar graph of fEPSP slope changes during FSK-IBMX in the presence of SKF38393 at 30PND and 90 PND MK-801 groups. **F)** Bar graph of fEPSP slope changes at 120min in presence of SKF38393, at 30 PND and 90 PND MK-801 groups. **G)** Bar graph of fEPSP slope changes at 240min in the presence of SKF38393, at 30 PND at 90 PND MK-801 groups.

When analyzing the last 5min of FSK-IBMX and SKF38393 perfusion, the fEPSP slope was above the baseline at 30 and 90 PND (at 30 PND, t=120.7, p<0.001; at 90 PND, t=40.80, p<0.001; Figure 5E). At 120min post-FSK-IBMX and SKF38393 perfusion, the fEPSP slope maintained above the baseline with a robust late-LTP (at 30 PND, t= 23.03, p<0.001; at 90 PND, t= 31.34, p<0.001; Figure 5F). Finally, at 240min after FSK-IBMX with SKF38393 perfusion, we observed a consistent increase in the fEPSP slope above the baseline levels at both 30 and 90 PND (at 30 PND, t= 6.81, p<0.001; at 90 PND, t= 58.19, p<0.001; Figure 5G). These findings show that activation of D1-like receptors during FSK-IBMX perfusion restores late-LTP expression in acute hippocampal slices of adolescent and young adult rats from the MK-801 schizophrenia-like model.

### 3.5. SKF38393 in the dorsal hippocampal CA1 area ameliorates retention phase impairment at the adolescent and adulthood stages in MK-801-treated animals in the Barnes maze test

To determine if activation of D1-like receptors can restore cognitive function in the MK-801 schizophrenia-like model, which was observed as an improvement of the Barnes maze retention phase parameters, we investigated the effect of SKF38393 or saline only into dorsal CA1 on memory retrieval in the Barnes maze task (see methods and experimental design, Figure 1, panel E, and schematic in Figure 3, panel A for details). MK-801 animals were divided into two groups and trained in the same conditions during the acquisition phase. At 30 PND, no statistical differences were observed between groups (on Day 1, latency: t=0.62, p=0.54, errors: t=1.30, p= 0.20, ns; or at Day 4, latency: t=2.0, p=0.06, errors: t= 1.56, p=0.13), thus evaluating them together, we observed a decreased latency and number of errors to find the target and enter into the goal chamber on day 4 compared to day 1 (for latency, t=7.059, p<0.001; for number of errors, t=5.46, p<0.001; Figure 6A-B). Similar to before, at 90 PND, no statistical differences were observed between groups (at Day 1, latency: t=0.95, p=0.34, errors: t=1.19, p= 0.24, ns; or at Day 4, latency: t=0.33, p=0.73, errors: t= 0.98, p=0.33, ns). When evaluating together, we don’t observe a decreased latency (t=1.91, p=0.06, ns), whereas the number of errors was reduced (t=3.03, p<0.01) when comparing day 4 with 1 (Figure 6C-D, respectively).

**Figure 6.**
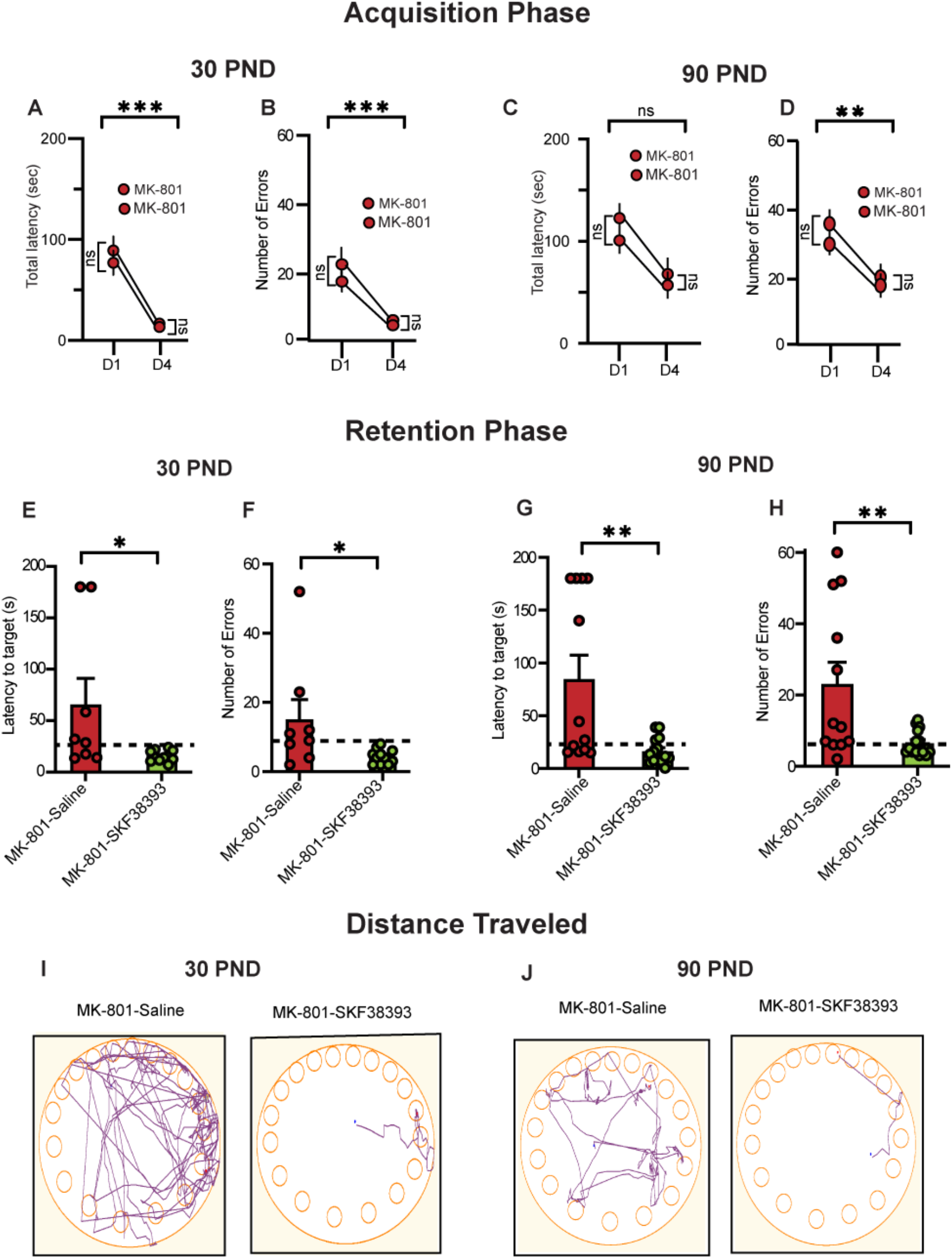
SKF38393 (D1-like agonist) infusion in dorsal CA1 ameliorates memory retrieval impairments in the Barnes maze retention phase in MK-801-treated animals at 30 and 90 PND. **A)** Habituation phase at 30 PND, latency to target on 1^st^ and 4^th^ day MK-801 groups. **B)** Habituation phase at 30 PND, number of errors on 1^st^ day and 4^th^ day MK-801 groups. **C)** Habituation phase at 90 PND, latency to target on 1^st^ day and 4^th^ day MK-801 groups. **D)** Habituation phase at 90 PND, number of errors on 1^st^ day and 4^th^ day MK-801 groups. **E)** Retention phase at 30 PND, latency to target on 5^th^ day in MK-801-saline (dark red symbols) and MK-801-SKF38393 (green symbols) groups. **F)** Retention phase at 30 PND, number of errors on 5^th^ day in MK-801-saline (dark red symbols) and MK-801-SKF38393 (green symbols). **G)** Retention phase at 90 PND, latency to target on 5^th^ day in MK-801-saline (dark red symbols) and MK-801-SKF38393 (green symbols). **H)** Retention phase at 90 PND, number of errors on 5^th^ day in MK-801-saline (dark red symbols) and MK-801-SKF38393 (green symbols). **I)** At 30 PND, distance traveled on the 5^th^ day in the MK-801-saline group and MK-801-SKF38393 group. **J)** At 90 PND, distance traveled on the 5^th^ day in MK-801-saline group and MK-801-SKF38393.

Furthermore, when evaluating the retention phase, groups were divided by age (30 and 90 PND) and by infusion of either saline (MK-801-saline) or SKF38393 (MK-801-SKF38393) into dorsal CA1. At 30 PND, we consistently observed a decreased latency and errors to find the escape hole (t=2.307, p=0.0339 and t=2.205, p=0.0415, respectively) compared to the MK-801-saline only group (Figure 6E-F). At 90, PND infusion of SKF38393 also ameliorates the impaired retention phase performance of MK-801 treated animals. We found that the MK-801-SKF38393 group has a decreased latency (t= 3.233, P=0.035; Figure 6G) and fewer errors (t= 2.881, p= 0.0082; Figure 6H) to locate the target hole and go into the goal chamber when compared to MK-801 saline group. Lastly, we tracked the distance traveled and observed that at 30 and 90 PND, MK-801-SKF38393 reduced the distance traveled to find the escape box compared to the MK-801-saline group (Figure 6I-J).

Taken together, these data show that activation of D1-like receptors with the agonist SKF38393 in dorsal CA1 hippocampal area can improve spatial retention deficits in animals neonatally treated with MK-801 during adolescent and adulthood stages that mimic the onset of schizophrenia-like symptoms.

## 4. Discussion

In the present study, we investigate the effects of MK-801 neonatally injected treatment on electrophysiological and behavioral changes during two developmental periods: adolescence and young adulthood. This neonatal treatment with an NMDA antagonist is considered an animal model of schizophrenia [49], [50], [51], and the two developmental stages studied are the prodrome phase (adolescence) and the time when schizophrenic symptoms usually appear (young adulthood) [3], [4], [6]. We found that both late-LTP and retention phase in the Barnes maze were impaired in MK-801-treated animals at both ages. Lastly, we observed that dopamine through D1-like receptor activation could ameliorate and restore synaptic plasticity and memory retention at both ages, revealing a new role for dopamine D1-like receptors in memory consolidation and retrieval associated with schizophrenia.

### 4.1 Late-LTP synaptic plasticity impairment in MK-801 neonatal schizophrenia-like model

*In vitro* studies suggest that consolidation of new synapses associated with late-LTP can lead to the synthesis of new proteins and mnemonic consolidation processes. A decrease or impairment in this phenomenon is associated with a malfunction during aging and disease course [52], [53]. A mixture of forskolin (FSK) and 3-isobutyl-1-methyl-xanthine (IBMX) is frequently used to induce a chemical type of late-LTP that can last for several hours and is dependent on NMDA receptors and phosphorylation of PKA kinases. Some of the advantages of chemical late-LTP induction are to activate the machinery of protein synthesis involved directly, maximizing the increase of the number of synapses that undergo the late-LTP and recruiting most of the cells into our brain slice preparation [32], [33], [34]. With this idea, our data indicates that late-LTP in control conditions is induced in hippocampal area CA1 from adolescent and young adult rats when a mixture of the FSK-IBMX bath perfusion was made; in both conditions, the magnitude of fEPSP slope increase was similar. Nevertheless, the same protocol could not induce a late LTP in the MK-801-treated animals.

Previous studies have described impairments in early-LTP phases in MK-801 schizophrenia-like model [27], [43] and during critical developmental stages, NMDA antagonists modify receptor subunits and union properties of ligands [9], [51], [54], modifying LTP induction and expression [28], [49]. Consistent with this idea, dysregulation in proteins underlying NMDA receptor activation in schizophrenia patients and animal models, including kinases and phosphatases, has been observed [55]. Thus, neonatal MK-801 treatment may cause an imbalance in intracellular calcium, leading to decreased activation of kinases and phosphatases that regulate synaptic plasticity mechanisms. In line with this idea, we previously reported that neonatal MK-801 treatment impairs early LTP since early adolescence, preceding the schizophrenia-like behavioral changes observed in early adulthood [27]. This data correlates with other studies that reported an impaired induction of early LTP at the adolescent and young adulthood stages, accompanied by cognitive impairment performance in the Morris water maze task [28], [43].

### 4.2 The retention phase of episodic memory associated with the CA1 area function is impaired in the MK-801 schizophrenia-like model

Episodic memory encoding and retention correlate with a precise hippocampal function [56], [57], [58]. We observed that adolescent and young adult rats have a consistent learning curve in control conditions, leading to robust retention during both age periods. Additionally, we observed that neonatal MK-801 treatment leads to increased latency and errors in the retention phase during adolescence. In this sense, the retention phase relies on proper hippocampal functioning, which involves recollecting previously acquired and consolidated memories [59], [60]. Studies indicate that in schizophrenia-like models, as the use of MK-801 treatment during the neonatal period or a prenatal immune activation, the decline in cognitive task performance during adolescence (40 PND) is significantly higher compared to control groups. This includes different behavioral tasks such as the object recognition in a novel contextual environment, the Morris water maze, and the Barnes maze [61]. Our data suggests that cognitive mechanisms related to hippocampal function in schizophrenia-like models begin to display impairments in early adolescence, with an impaired performance during the Barnes maze retention phase. Interestingly, during this period, no other schizophrenia-like behaviors were seen as an increased response in the pre-pulse inhibition test, and diminished social behaviors were observed [27].

Additionally, we observed consistent impairment during young adulthood in the Barnes maze retention phase. Similar to our data, other studies indicated how MK-801 neonatal treatment induces impairment in the Morris water maze [4], the Barnes maze [29], and the radial maze [50]. Other schizophrenia-like models reported similar effects in the Y-maze passive avoidance and Barnes maze during young adulthood [61], [62].

Cognitive deficits in young adults neonatally treated with MK-801 are accompanied by behaviors that resemble negative and positive symptoms of schizophrenia. Our previous report shows that treating neonates with MK-801 leads to impaired sensorimotor processing and decreased sociability during the social interaction test [27]. Additionally, Uehara et al. reported an increase in the startle response index in the pre-pulse inhibition test and hyperlocomotion during young adulthood [63]. Our data suggests that a disruption in critical stages of neurodevelopment leads to cognitive deficits during young adulthood that mimic symptoms of schizophrenia.

### 4.3 Dopamine neuromodulation through activation of D1-like receptors as a rescue mechanism in the MK-801 schizophrenia-like model

The consolidation and retrieval of cognitive processes is correlated with late-LTP expression that requires dopamine as a neuromodulator [12], [64], [65], [66], [67]. *In vitro* studies have indicated that modulation of dopamine through D1-like receptors supports the induction and maintenance of late-LTP for hours [12], [67]. Meanwhile, an aberrant modulation of dopamine through the inactivation of dopamine D1-like receptors impairs the induction of late-LTP [18], [68], [69]. In line with this, we observed that blockade of D1-like receptors with SCH23390 antagonist in control conditions leads to an aberrant performance in the retrieval stage of the Barnes maze in juvenile and young adolescent rats. Silva et al. (2012) reported that blocking D1-like receptors in the dorsal CA1 area leads to increased parameters in the Morris water maze during the retrieval stage [70]. Previous studies reported that blocking D1-like, but not D2-like, receptors impairs late-LTP, highlighting the essential role of D1-like receptor activation in memory retrieval in the hippocampal CA1 area [71], [72]. Furthermore, we observed that activation of D1-like receptors with SKF38393 in the MK-801 treated animals reestablished late-LTP and maintained potentiated for several hours. Previously, our research group reported that a D1-like agonist could partially restore the plasticity capabilities of pyramidal neurons in CA1 [31]. To our knowledge, these are the only studies investigating the effect of D1-like agonists on restoring cellular and behavioral impairments in the hippocampal CA1 area in MK-801 neonatal-treated animals.

The dopamine circuit, through D1-like receptor activity, has gained interest in the neurodegeneration of Alzheimer’s disease. Xiang et al. (2016) observed that a D1-like activation can protect late-LTP of the hippocampal CA1 region of oligomeric amyloid-beta deposits [73]. Thus, dopamine neuron loss may contribute to memory and reward dysfunction in mouse models of Alzheimer’s disease and other brain conditions [73], [74]. Together, these results reveal that dopamine, through D1-like receptor activation, helps improve memory consolidation and retrieval in different neurological disorders.

One goal of scientific research is to uncover different pathways that lead to treatment to lessen or prevent schizophrenia symptoms. In this line, other studies have focused on preventing the manifestation of cognitive deficits with either an enriched environment or the treatment with herbal alkaloids during different developmental stages in the MK-801 schizophrenia model [75], [76]. Although these studies provide insight into treating schizophrenia symptoms, this leads to the question of a potential treatment once the neonatal disturbances have occurred. Schizophrenic patients receive pharmacological treatment to block dopaminergic transmission and decrease positive and negative symptoms, with a mild or null effect on cognitive symptoms [77]. During schizophrenia, dopaminergic levels decreased in the mesocortical region with a decreased expression in D1-like receptors, indicating dopamine’s role in cognitive symptoms [21], [78]. In line with this idea, we observed that infusions of dopamine D1-like agonist (SKF38393) in hippocampal area CA1 during adolescence and young adulthood improved impairments in Barnes maze performance and late-LTP expression in the MK-801 model, suggesting a new role of dopaminergic modulation in cognitive impairments of schizophrenia models. In physiological conditions, dopamine plays a crucial role in episodic memory formation, consolidation, and retrieval in the hippocampal CA1 region [20], [79]. In the passive avoidance test, Broussard et al. (2016) demonstrated that dopaminergic D1-like activity is necessary to retain aversive memory [20]. Furthermore, McNamara et al. reported that activating the locus coeruleus projections into the hippocampal CA1 area through optogenetics can enhance novel object exploration and maintain cognitive map stability [25]. Additionally, Xiang et al. (2010) showed that D1-like but not D2-like activation is critical for spatial learning in the Morris water maze [80]. Melief et al. (2015) found that activating the dopaminergic D1 receptor with SKF38393 agonist improved the Barnes maze performance during the retention phase in a mouse model with Alzheimer’s disease [81]. In line with these reports, we discovered that administering a dopaminergic D1-like agonist (SKF38393) locally in the dorsal CA1 can reduce cognitive deficits associated with memory retrieval in MK-801 neonatal-treated animals, assessed during childhood and young adulthood. This result correlates with previously reported data showing that SKF38393 treatment in a genetic schizophrenia mouse model improved performance in the Y-maze [82]. The present study supports that D1-like receptor activation improves cognitive performance during memory retrieval once MK-801 neonatal treatment has disturbed physiological neurodevelopmental processes.

## 5. Conclusion

Our results suggest that episodic memory linked to hippocampal function is impaired in the MK-801 schizophrenia-like model, provoking an early onset of memory retrieval impairments associated with hippocampal area CA1 that is long-lasting from adolescence until young adulthood, imitating the clinical symptoms of schizophrenia disease.

Our data also revealed that dopamine circuit activation through D1-like receptors in the MK-801 model promotes the induction of synaptic plasticity and cognitive performance similar to healthy conditions. These results indicate an important role of dopamine dysregulation in hippocampal area CA1 in schizophrenia-like symptoms with a possible key role in the rescue of cognitive functions.

## Acknowledgments

This work was supported by a grant from the Consejo Nacional de Ciencia yTecnología (CONACYT), Ph.D. fellowship 281099 to MH-F. We thank Drs. Carmen Vivar and Jayeeta Basu for feedback and comments on the manuscript, and Cara D. Johnson for help proofreading. We also thank Juan Javier López Guerrero and Isabel Beltran Villalobos for their technical assistance.

## Author Contributions

MH-F conceptualized the study, performed the experiments and data analysis, and wrote the original draft. CL-R conceptualized the study, provided resources, supervised the work, and prepared the final version of the manuscript. EJG provided resources and participated in the experimental design.

